# Temperature and Ultraviolet Radiation Drive Divergent Visible and Near-Infrared Reflectance Patterns in Butterflies

**DOI:** 10.64898/2025.12.16.694481

**Authors:** Calista Z.E. Kou, Changku Kang, Matilda Brindle, Ilya M. D. Maclean, Jon Bridle, Alex L. Pigot, Robert J. Wilson, Joseph Williamson

## Abstract

1. Climate change poses extreme risks to biodiversity, threatening the ecosystems upon which humanity depends. Understanding the traits that mediate how organisms respond to climatic gradients in space and time will enable better predictions of the taxa and ecological communities most at risk from warming.
2. In insects, colouration influences thermoregulation, but warming responses must be traded off against other selective pressures. We hypothesise that butterfly wing reflectance in visible and near-infrared (NIR) spectra respond differently to temperature and ultraviolet radiation over space and time. We combined existing reflectance data of 97 butterfly species in both NIR and visible wavelengths, with long-term abundance monitoring data of 120 butterfly communities sampled across 1650 m altitude gradients from May to August in two time periods (2004/5 and 2017).
3. The visible and NIR reflectances of butterfly communities showed a non-linear relationship with altitude, with the lowest reflectance (darkest butterflies) at the coolest, highest sites. In contrast, community visible reflectance decreased through the year as temperatures warmed over spring-summer, whereas community NIR reflectance remained constant, revealing divergent responses of reflectance types to seasonal changes. Temperature had opposing effects on visible and NIR reflectance of butterfly communities, where increasing temperature reduced community visible reflectance strongly while increasing community NIR reflectance slightly.
4. Considering the effects on reflectance of shared evolutionary history in a Bayesian hierarchical model for individual species, lighter-coloured (more reflective) species were associated with warmer temperatures - flying later in spring-summer or at lower altitudes. However, instead of decreasing in reflectance through the year and across temperature gradients, species instead became lighter, we expect as a result of a Simpson’s paradox.
5. These results emphasise how visible and NIR reflectance wavelength bands mediate butterflies’ responses to environmental gradients in distinct ways, despite being highly correlated across species. We also show that incorporating phylogeny into trait-environment models is essential; relying on traits alone would lead to incorrect inferences and predictions of taxa most at risk from warming climates.

## 1 Introduction

Anthropogenic atmospheric carbon emissions are driving rapid changes in global temperatures, posing critical risks to biodiversity (Calosi *et al*., 2008; Bellard *et al*., 2012; Illán *et al*., 2012; Urban, 2015). Species are already exposed to elevated thermal stress, driving range shifts (Wilson *et al*., 2005; Bellard *et al*., 2012; Wiens, 2016), population declines and local extinctions as habitats become unsuitable (Calosi *et al*., 2008; Wiens, 2016; Román-Palacios and Wiens, 2020). These risks for biodiversity will increase over the coming century (Parmesan *et al*., 2022), threatening the productivity of ecosystems and the stability of the biosphere on which humanity depends (Rockstrom *et al*., 2009; Bridle *et al*., 2025). Understanding how organisms respond to climatic gradients in both time and space, and which traits mediate said responses, will help us to predict taxa and systems most at risk from rising temperature (Chevin and Bridle, 2025; Williamson *et al*., 2025).

Traits that relate directly to the thermal biology of phenotypes are key candidates for predicting species’ responses to warming. Although historically, physiological, mostly stress-resistance traits have dominated assessments of vulnerability to warming (Deutsch *et al*., 2008; Angilletta Jr, 2009; Pinsky *et al*., 2019), traits that mediate the exposure of organisms to temperature in advance of such stress responses are also important (Briscoe *et al*., 2023; Pottier *et al*., 2025). Colouration is determined by the intensity and wavelength of light organisms reflect, which in turn affects heat absorption from radiation (Medina *et al*., 2018; Kang *et al*., 2021) and thus the operative body temperature of organisms in the wild (Stuart-Fox *et al*., 2017). Understanding how reflectance will mediate the effects of rising temperatures may be essential for predicting how species respond to climate change.

Organismal colouration varies dramatically across temperature gradients (Bishop *et al*., 2016; Gautam and Kunte, 2020; Law *et al*., 2020). In general, darker colours, which drive higher operative body temperature through greater absorbance of environmental radiation, are favoured in cooler climates (Bogert, 1949). For example, ant, butterfly and dragonfly assemblages are darker in cooler regions (Zeuss *et al*., 2014; Bishop *et al*., 2016; Kang *et al*., 2021). In addition to geographic patterns, South African ant assemblages (Bishop *et al*., 2016) and European dragonfly assemblages (Zeuss *et al*., 2014) have become lighter on average as the climate has warmed in recent decades.

However, colouration is not just shaped by the selective pressures of the thermal environment (Stuart-Fox *et al*., 2017). Organisms use colouration for camouflage (Kettlewell, 1955), social signalling (Wiernasz, 1995), and protection from ultraviolet (UV) radiation (Majerus, 1998), desiccation (Kalmus, 1941; Parkash *et al*., 2008; Parkash *et al*., 2009; King and Sinclair, 2015) and parasites (True, 2003; Cuthill *et al*., 2017; Schirmer *et al*., 2023). Given these pressures can select for phenotypes that do not maximise thermal tolerance, darker organisms do not always dominate cooler climates. For example, ant assemblages are darker in the hotter forest canopies compared to the forest floor (Law *et al*., 2020), while dung beetle communities initially become lighter as forest air temperatures increase, before getting darker again (Williamson *et al*., 2022).

Although previous studies have investigated how colouration mediates thermoregulation in ectotherms, most have focused on reflectance within the visible spectrum (400 – 700 nm) (Stuart-Fox *et al*., 2017; Munro *et al*., 2019; Kang *et al*., 2021). However, near-infrared (NIR) wavelengths (700 – 2500 nm) account for approximately 55 % of the energy in solar radiation (Stuart-Fox *et al*., 2017), meaning that NIR radiation is an often-overlooked but critical driver of organismal body temperature. In contrast to visible wavelengths, NIR reflectance is invisible to most organisms and thus less likely to be shaped by selection pressures such as social signalling (Luo *et al*., 2011; Stuart-Fox *et al*., 2017). At higher altitudes, temperature declines while UV radiation increases (Figure 1d), meaning that darker species tend to dominate these environments (Reguera *et al*., 2014; de Souza *et al*., 2020) because they benefit from both higher UV protection and body temperature. However, temperature and UV do not always negatively covary. Temperatures decrease and UV increases towards the poles, and several recent studies have shown that the relationship between climate and reflectance was stronger for NIR than visible wavelengths through space over latitudinal gradients (Medina *et al*., 2018; Munro *et al*., 2019; Kang *et al*., 2021). Similarly to their relationship over latitudinal clines, UV and temperature both peak during summer in temperate zones (Figure 1e), though it remains to be determined how this influences organismal reflectance.

**FIGURE 1.**
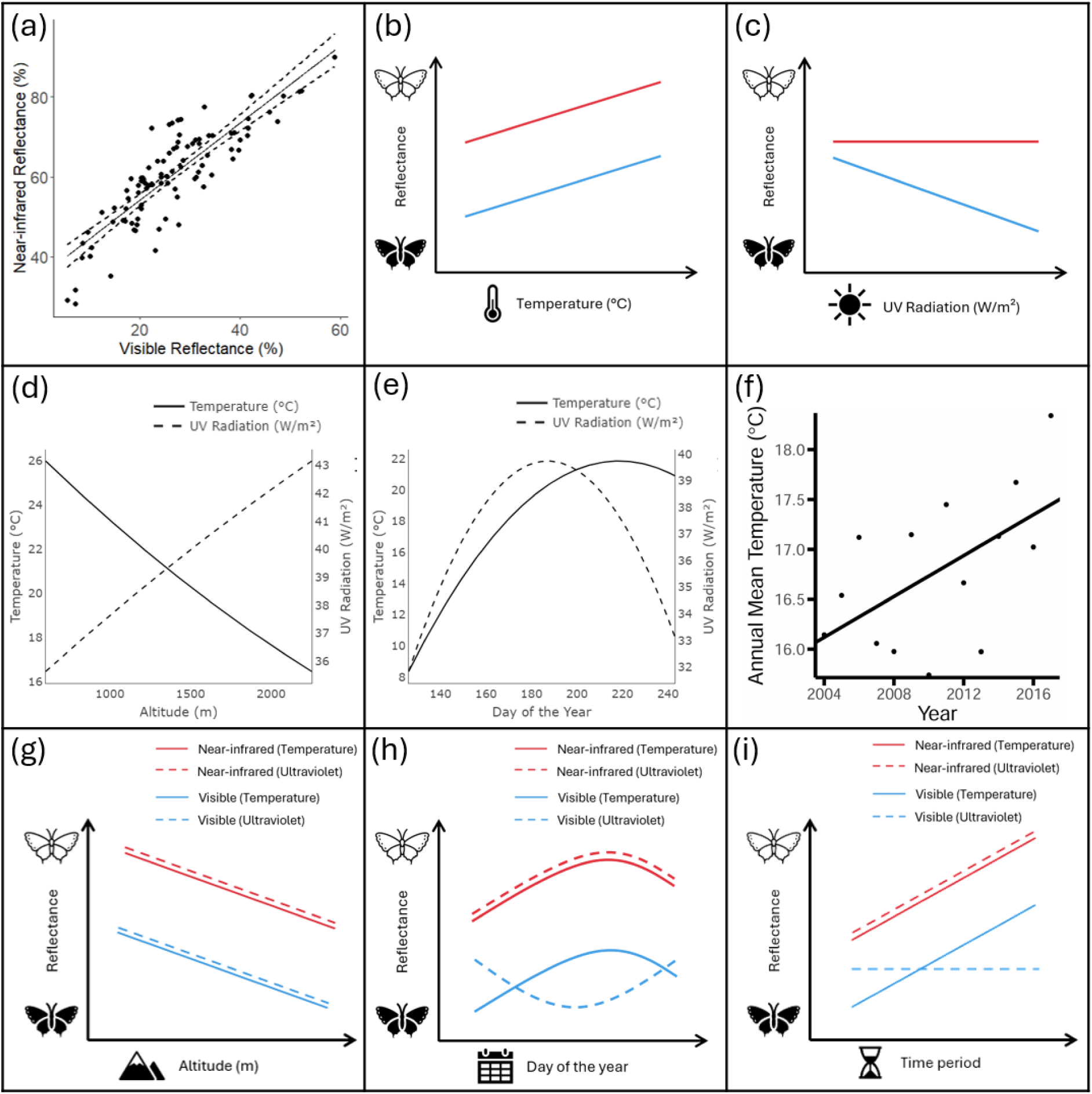
An infographic depicting our hypothesised relationships between butterfly reflectance and climatic and spatiotemporal variables. (a) Scatterplot showing the relationship between visible and near-infrared (NIR) reflectance of butterfly species. (b) Hypothesised relationship between visible (blue) and NIR (red) reflectances and temperature. (c) Hypothesised relationship between visible (blue) and NIR (red) reflectances and ultraviolet (UV) radiation. (d) Relationship between altitude and mean annual temperature (solid line) and UV radiation in our study system (dashed line). Mean annual temperature was downscaled from coarse-scale ERA5 climate data using the *mesoclim* package (Maclean and Mosedale, 2025) and UV was modelled using *NicheMapR (ref)* (see methods for details). (e) Relationship between day of the year and mean temperature of the past 30 days (solid line) and mean daily total UV radiation of the past 30 days (dashed line). (f) Annual mean temperatures over the course of study period in the region. (g–i) Hypothesised relationships between visible (blue) and NIR (red) reflectance and (g) altitude, (h) day of the year and (i) time period due to both temperature (solid line) and UV radiation (dashed line).

To test how visible and NIR reflectance patterns vary over climatic gradients in space and time, we exploited an existing dataset of visible and NIR reflectance of butterfly species measured from museum specimens (Kang *et al*., 2021), combined with butterfly abundance data from long-term monitoring in the Sierra de Guadarrama in Central Spain (Wilson *et al*., 2007; Álvarez *et al*., 2024; Goded *et al*., 2024). Butterflies exhibit diverse wing colouration (Beldade and Brakefield, 2002) and rely on external mechanisms to regulate their body temperature, making them highly sensitive to temperature changes (Kingsolver, 1985a). Effective thermoregulation is vital for butterfly survival, enabling essential activities like flight (Kingsolver, 1988) and preventing heat death (Tsai *et al*., 2020). As with patterns observed in other taxa, butterfly colouration varies across climatic gradients, with darker species tending to occur in cooler climates and lighter species in warmer climates (Xing *et al*., 2016; Kang *et al*., 2021). The Sierra de Guadarrama mountains exhibit strong temperature and UV gradients in both space and time while hosting high butterfly species richness (> 100 species) (Wilson *et al*., 2005; Wilson *et al*., 2007; Mingarro *et al*., 2021), making this study location particularly suitable for studying variation in visible and NIR reflectance patterns. We hypothesised that as temperature increases, both visible and NIR reflectance of butterfly communities and species will increase, as darker colouration allows butterflies to prevent overheating at high temperatures (Figure 1b). At higher UV radiation, we predict that while butterfly communities and species will have lower visible reflectance to protect biomolecules from UV radiation (Figure 1c).

To understand how observed patterns of reflectance vary across spatiotemporal gradients, we also examine how community-level and species-level visible and NIR reflectance of butterflies varies across altitude, through days of the year when butterflies are active, and across our two sampling time periods (2004/5 and 2017). We predict that visible and NIR reflectance will decrease as altitude increases, due to lower temperatures and higher UV radiation at higher altitudes (Figure 1g). Through the year, we expected NIR reflectance to increase as temperatures rise, but for visible reflectance to either increase with elevated temperatures, or decrease with higher UV radiation (Figure 1h). We also predicted that both reflectance types will increase over the years due to ongoing warming in the region (Figure 1i). To test these hypotheses, we generated weighted-mean reflectance traits for communities recorded on specific dates at specific sites and related these to our spatiotemporal and climatic factors. To account for the potentially confounding effects of shared evolutionary history on traits, we also used phylogenetic species-level models to predict trait values using species’ spatiotemporal and climatic associations.

## 2 Materials and Methods

### 2.1 Butterfly abundance data

Butterfly abundances for 97 species were surveyed at 120 transects in the Sierra de Guadarrama mountain range in Central Spain (4°32’–3°39’ W, 40°34’–41°25’ N) from May to August, 4-5 times at each transect in our two time periods (2004/5 and 2017). For each survey, butterflies were identified and counted in a 5 x 5 m virtual sampling box in front of the recorder along 500m transects walked at a constant speed every two to three weeks (also known as a Pollard Walk) (Pollard and Yates, 1993; Illán *et al*., 2010). Species from two genera (*Carcharodus* and *Pyrgus*) were not included in this study due to difficulties with identification *in situ*.

### 2.2 Reflectance data

Reflectance data were taken from those generated by Kang *et al*. (2021), who measured both visible and near-infrared (NIR) reflectance of butterfly specimens from the Natural History Museum in London. Briefly, two specimens were photographed for each species using a spectrum converted DSLR camera positioned below two light bulbs to prevent light from shining directly into the camera lens. Reflectance spectra were captured using a camera fitted with lens filters that completely blocked wavelengths outside the transmission range to capture visible (400 – 680 nm) and NIR (670 – 1050 nm) wavelengths. Mean reflectance values were measured using ImageJ for six body regions: dorsal thorax, dorsal basal wings, dorsal entire wings, ventral thorax, ventral basal wings and ventral entire wings. RGB values were scaled linearly, resulting in reflectance values of 0 – 100 %. The mean reflectance of all six body regions was calculated to give the average reflectance of each species. For our community-level analysis, community-weighted means (CWMs) of the average visible and NIR reflectance were calculated for each survey using the formula:

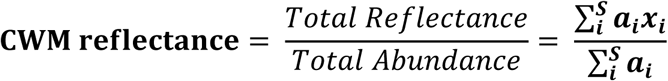

Where 𝑺 is the species richness, 𝒂_𝒊_is the abundance of a species and 𝒙_𝒊_is the average (visible or NIR) reflectance of a species.

### 2.3 Climate data

To estimate the temperatures that our studied communities are exposed to, we generated hourly mesoclimate data for the 30 days before each survey by downscaling coarse-scale climate data from the ERA5 database (Hersbach *et al*., 2020) using the *mcera5* package (Klinges *et al*., 2022). To do so, we downloaded a digital terrain model at 6.48 x 10-4 ° (or ∼72 m latitude x ∼54 m longitude) resolution was accessed from the AWS Terrain Tile Service (https://registry.opendata.aws/terrain-tiles/) using the elevatr package (Hollister *et al*., 2023) in a 200 m buffer around the transect centroid. Regional climate data were then downscaled using the standard equations available in the *mesoclim* package (Maclean and Mosedale, 2025). For each survey, the mean temperature of the preceding 30 days was taken across the 200 x 200 m grid.

We generated clear-sky ultraviolet (UV) radiation values at each transect location by inputting the longitude, latitude and altitude of each transect centroid into the “micro_global” function from the *NicheMapR* R package (Kearney and Porter, 2020). This function uses a high-resolution global climate dataset (New *et al*., 2002) to calculate solar irradiance across the full solar spectrum, including UV wavelengths (280 – 400 nm). We simulated scattered solar radiation, adjusted for terrain effects and produced wavelength-specific solar radiation. For each survey, we summed the hourly UV radiation values to calculate the total daily UV radiation, then averaged the daily totals over the 30 days prior to sampling to obtain the mean daily UV radiation for each survey.

### 2.4 Statistical analyses

#### 2.4.1 Community-level reflectance responses to spatiotemporal and climatic gradients

To estimate how community-weighted mean reflectance varies across space and time, we fit a linear mixed-effects model in which community reflectance was predicted by a second-order polynomial of altitude, day of the year and time period (2004-2005 or 2017), together with their interactions with reflectance type (NIR or visible). Transect was included as a random effect, with date nested within transect to account for non-independence arising from repeated measurements of NIR and visible community reflectances from the same survey.

To test whether community reflectance varies over climatic gradients, we first fit a linear mixed-effects model of community-weighted mean reflectance as predicted by reflectance type and mean temperature of the past 30 days as fixed effects. Due to covariance between temperature and UV, we fit a separate model of community-weighted mean reflectance as predicted by reflectance type and mean UV radiation of the past 30 days, and their interaction. In both the temperature and UV model, transect and date nested within transect were included as random effects.

All spatiotemporal and climatic mixed-effects models were selected using the *lme4* package (Bates *et al*., 2014). Best fitting models were selected using likelihood ratio tests (LRT) to sequentially remove non-significant terms from the full models.

#### 2.4.2 Species-level reflectance responses to spatiotemporal and climatic gradients

To control for the effects of phylogeny on the species-level response of traits to spatiotemporal and climatic gradients, we used a publicly available time-scaled, multi-gene phylogenetic tree (Wiemers *et al*., 2020). We converted two artefactual negative branch lengths, within *Melitaea* and *Hipparchia*, to 0 (Wiemers, personal communication), and then pruned the tree to include only the 97 butterfly species present in our surveys.

Phylogenetic signal of butterfly visible and NIR reflectance were measured using Pagel’s lambda (λ), which measures the extent to which closely related species express similar phenotypes. The statistical significance of λ was evaluated using LRT against models of random evolution.

To estimate species-level responses to spatiotemporal gradients, we generated abundance-weighted mean flight date and abundance-weighted mean altitude for each species. To capture species responses to climate change between our study periods, we calculated the difference between the mean abundance over two time periods (2004/5 and 2017) across all surveys. Similarly, for our climatic gradients, we used our metric of UV and temperature from the previous 30 days for each visit to a transect. We then took the abundance-weighted mean of these variables for each species, giving us a measure of each species’ thermal and UV radiation associations, hereafter referred to as abundance-weighted mean thermal and abundance-weighted mean UV association.

To test whether reflectance predicted species-level responses to spatiotemporal and climatic gradients, we fitted Bayesian hierarchical models using the *brms* R package (Bürkner, 2021). In our spatiotemporal models, species’ mean reflectance (visible or NIR) was our response variable, and our fixed effects were the species’ mean flight day, species’ mean altitude (both weighted by abundance) and the difference in mean abundance between our two time periods (2004/5 and 2017). In our climatic models, species’ mean reflectance (visible or NIR) was our response variable, and our fixed effect was either the mean temperature a species was recorded at, or species’ mean UV (both weighted by abundance), giving us a total of six models. All models contained a phylogenetic covariance matrix as a random effect. Models were run across four chains with a burn in of 5000 iterations, and a subsequent run of 45000 iterations with delta adapted to 0.99. Model trace plots and posterior distributions were visually inspected ensure good model fit. R-hat values were <1.01 and effective sample sizes were greater than 400 (100 times the number of chains). Statistical differences were considered strong or moderate if the 95 or 80 % Bayesian credible intervals (BCIs), respectively, did not cross zero (Brindle *et al*., 2023).

All analyses were performed using R Statistical Software v4.3.3 (R Core Team, 2024).

## 3 Results

A total of 59139 butterflies representing 97 species were recorded. Species reflectance varied broadly, with visible reflectance ranging from 8.3 - 46.6 % reflectance (mean = 26.1 % reflectance) and near-infrared (NIR) reflectance from 35.2 - 77.0 % reflectance (mean = 61.8 % reflectance). Visible and NIR reflectance were strongly correlated for species in our study (r = 0.85, p < 0.05; Figure 1a), which we expected as visible and NIR wavelengths are close together on the electromagnetic spectrum (Stuart-Fox *et al*., 2017). Therefore, it would be expected for both visible and NIR reflectance to show similar patterns across spatiotemporal and environmental gradients.

### 3.1 Community reflectance responses to spatiotemporal gradients

The best fitting model for our spatiotemporal analysis included interaction terms between reflectance type and altitude, day of the year and year category. Including a quadratic term for altitude significantly improved model fit (𝑥_2_^2^ = 15.01, p < 0.001, LRT compared to model with altitude as a linear term; Figure 2a, b). Community visible and NIR reflectance remained relatively flat at lower altitudes, before decreasing at higher altitudes (Figure 2a, b), suggesting a threshold response to altitude where butterfly communities become significantly darker after above a certain altitude.

**FIGURE 2.**
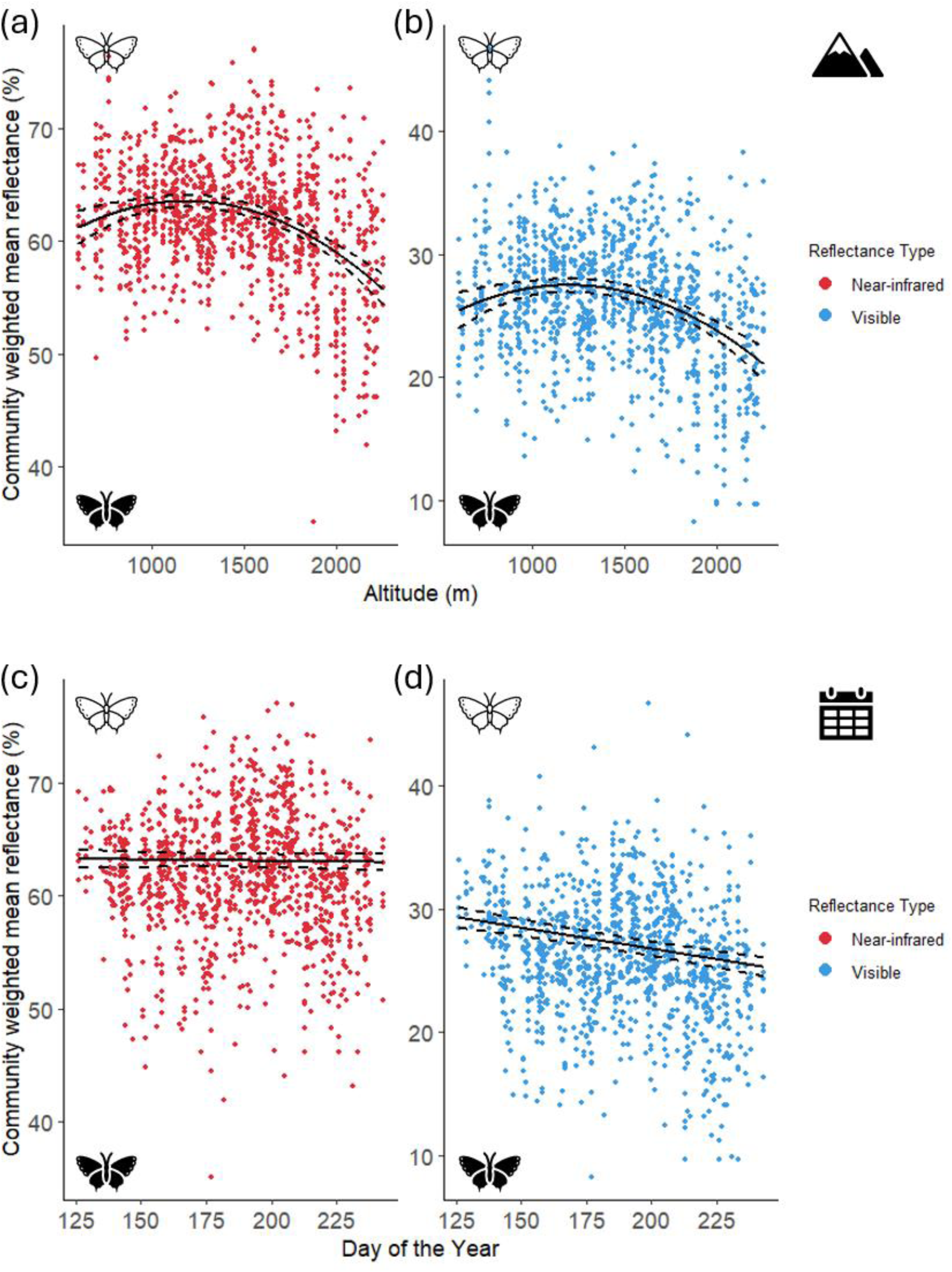
Scatterplots of the relationships between altitude and community-weighted mean (CWM) reflectance for (a) near-infrared (NIR) reflectance and (b) visible reflectance, and between day of the year and CWM reflectance for (c) NIR reflectance and (d) visible reflectance. Dashed lines denote 95 % confidence intervals.

Day of the year significantly improved model fit (𝑥_2_^2^ = 152.01, p < 0.001, LRT compared to model without day of the year; Figure 2c, d). Each later day in the year was associated with 0.032 % reflectance decrease in community visible reflectance (Figure 2d). However, community NIR reflectance remained constant throughout the year (Figure 2c). Additionally, species with higher NIR to visible reflectance ratios were more abundant later in the year (Figure S1).

Time period was not a significant predictor of either community visible or NIR reflectance.

### 3.2 Community reflectance responses to climatic gradients

Community-level reflectance was best predicted by a model including an interaction term between reflectance type and temperature ( 𝑥_1_^2^= 101.84, p < 0.001, LRT compared to the model without the interaction; Figure 3a, b). Community visible reflectance decreased strongly with increasing temperature (Figure 3b), where an increase in mean temperature of 1°C was associated with a 0.198 % reflectance decrease in community visible reflectance. In contrast, community NIR reflectance increased with increasing temperature (Figure 3a), where an increase in mean temperature of 1°C was associated with a modest 0.067 % reflectance increase in community NIR reflectance, despite the strong correlation of NIR reflectance with visible reflectance (Figure 1a).

**FIGURE 3.**
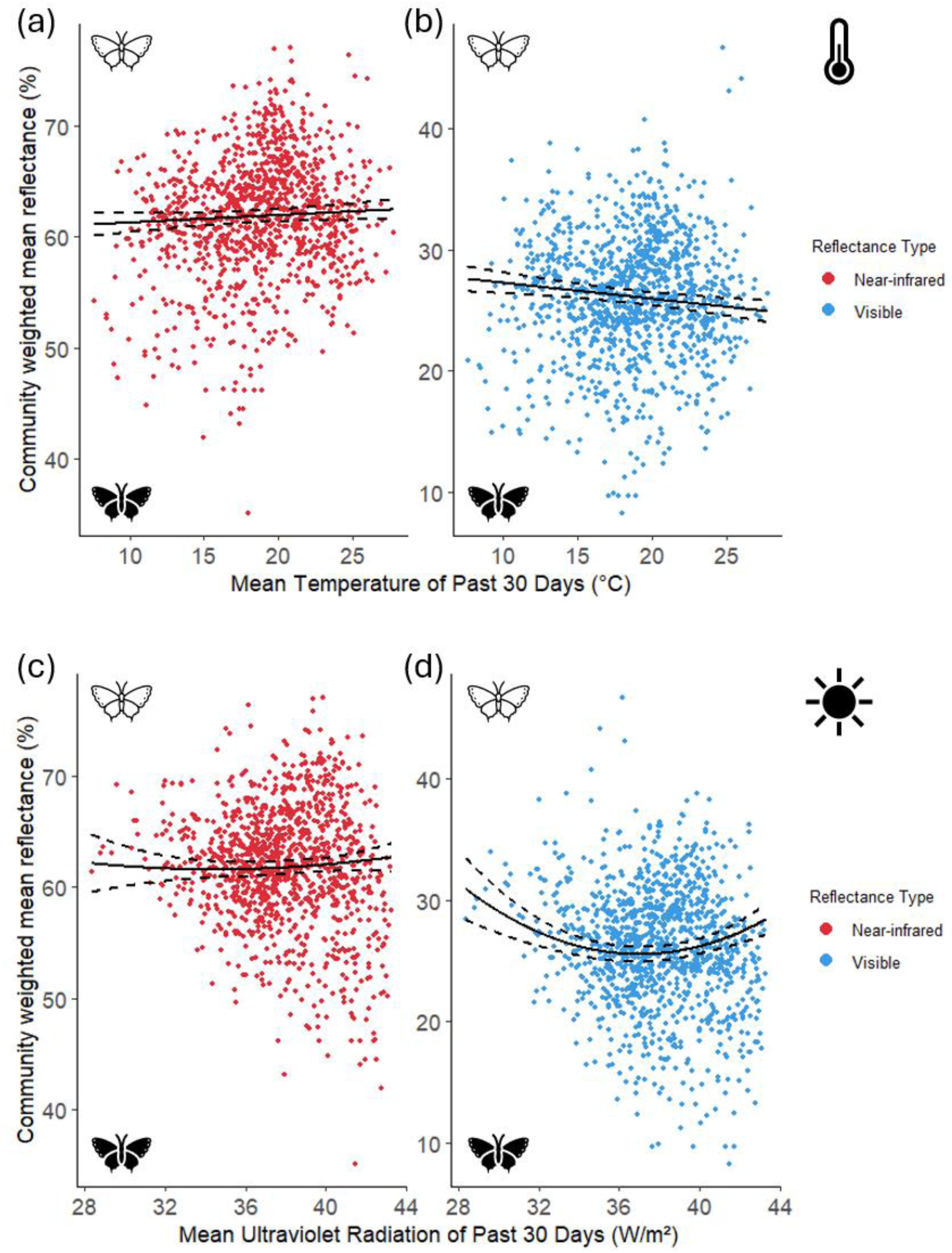
Scatterplots of the relationships between climatic variables and reflectance. Mean temperature and (a) community-weighted mean (CWM) near-infrared (NIR) reflectance and (b) CWM visible reflectance. Mean ultraviolet radiation and (c) CWM NIR reflectance and (d) CWM visible reflectance. Dashed lines denote 95 % confidence intervals.

Community-level reflectance was best predicted by a model including an interaction term between reflectance type (𝑥_2_^2^ = 44.08, p < 0.001, LRT compared to the model without the interaction; Figure 3c, d) and a quadratic effect of ultraviolet (UV) radiation (𝑥_2_^2^ = 50.07, p < 0.001, LRT compared to the model with UV radiation as a linear term; Figure 3c, d). Community visible reflectance decreased with increasing UV radiation, but increased slightly at very high UV levels (Figure 3d). In contrast, community NIR reflectance showed weaker, more linear changes across the UV gradient (Figure 3c). This indicates that butterfly communities became darker at intermediate UV levels for visible reflectance, while NIR reflectance remained relatively stable, despite strong correlation of NIR reflectance with visible reflectance (Figure 1a).

### 3.3 Species-level reflectance responses over spatiotemporal and climatic gradients

There was high phylogenetic signal in both visible (λ = 0.872, *p <* 0.001) and NIR (λ = 0.686, *p <* 0.001) reflectance, indicating that more closely-related species express similar reflectance phenotypes. Community-level results were corroborated by patterns of decreasing species-level reflectance with weighted-mean altitude in both visible (- 1.34 % per 1000 m; Table S1; Figure 4) and NIR (- 2.56 % per 1000 m; Table S1; Figure 4). Similarly, species-level NIR reflectance was not predicted by mean abundance shifts between the two sampling periods, though there was moderate support for butterflies with lower visible reflectance experiencing more positive abundance shifts from our first time period in 2004/5 to the second in 2017 (- 0.87 % per individual increase; Table S1; Figure 4). Contrastingly to our community-level patterns of reflectance that decreased through the year, species-level reflectance increased through the year with weighted-mean flight day in both visible (1.38 % per month; Table S1; Figure 4) and NIR wavelengths (2.66 % per month; Table S1; Figure 4), indicating that species are lighter later in the year once the strong effect of phylogeny is removed. Similarly, weighted-mean temperature of the previous 30 days was positively associated with reflectance in both visible (1.27 % per °C; Table S1; Figure 4) and NIR (2.52 % per °C; Table S1; Figure 4) wavelengths. Weighted-mean UV exposure of the previous 30 days was not associated with reflectance (Table S1; Figure 4).

**FIGURE 4.**
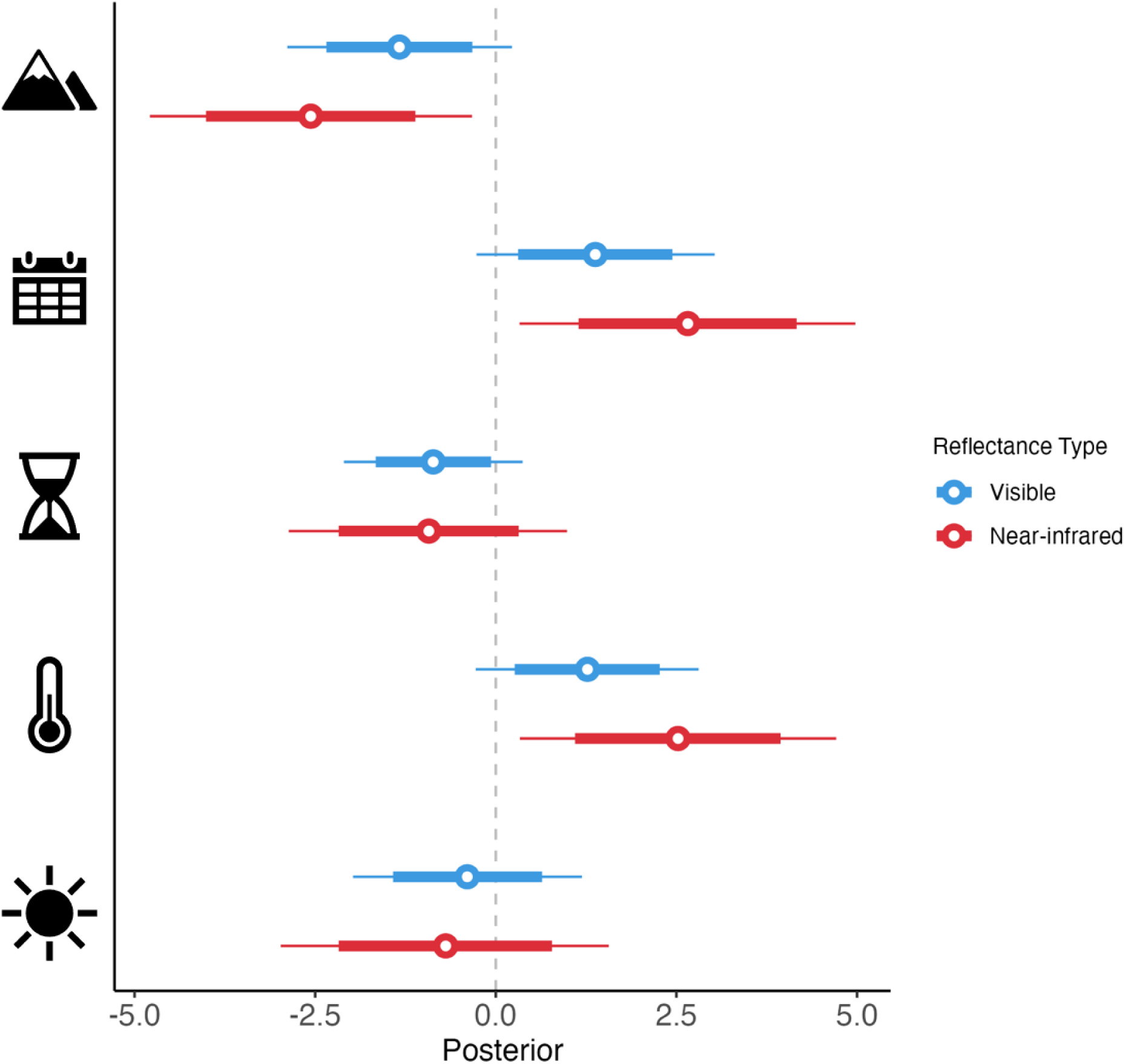
Standardized estimates for species-level models of reflectance. Estimates depicted (top to bottom): weighted-mean altitude, weighted-mean day of the year, mean abundance shift from 2004/5 to 2017, weighted-mean temperature, and weighted-mean UV. Hollow circles denote posterior medians, thick lines denote 80 % Bayesian credible intervals (BCIs), and thin lines denote 95 % BCIs.

## 4 Discussion

By combining abundance data from an altitude gradient in Central Spain (Goded *et al*., 2024) with species-specific reflectance data (Kang *et al*., 2021) and fine-scale climate models, we show that butterfly communities exhibit variation in colouration through both space and time, as well as temperature and ultraviolet (UV) radiation, through changes in the relative abundance of different species. Although the relationships between community colouration and our spatiotemporal gradients of altitude, day of the year and time period (2004/5 and 2017) generally support our initial hypotheses, the relationship of community-level and species-level reflectance with temperature and UV are complex. We suggest that this is due to contrasting relationships between UV and temperature in space and time, where UV and temperature are negatively correlated across space (with altitude) and positively correlated through time (day of the year). As reflectance in near-infrared (NIR) and visible wavelengths were also highly correlated across species, we expect that complex patterns may be a product of trade-offs of tolerances to divergent selective pressures. Our analyses reveal that visible and NIR reflectance respond differently across these gradients, highlighting the functional importance of NIR reflectance as a trait influencing species’ responses to temperature.

At higher altitudes, where temperatures are lower and UV radiation is higher, butterfly communities were darker. Similarly, butterfly species associated with higher elevations had darker phenotypes when controlling for the effect of shared evolutionary history. This pattern corroborates previous studies of colouration patterns across altitudinal gradients for multiple taxa, including butterflies, ants, wasps and lizards (Reguera *et al*., 2014; de Souza *et al*., 2020; Gautam and Kunte, 2020; Law *et al*., 2020; Kang *et al*., 2021), and supports the hypothesis that darker pigmentation is favoured in cooler environments to enhance heat gain from solar radiation (Bogert, 1949), and in higher UV environments to protect from molecular damage (Bastide *et al*., 2014; Delhey, 2017). We observed a polynomial relationship between reflectance and altitude, where butterfly reflectance remained relatively constant at lower altitudes before decreasing strongly as altitude near the tops of mountains. This non-linear relationship could be due to differences in habitat types across altitudes influencing vegetation cover (Illán *et al*., 2012; Mingarro *et al*., 2021), which can buffer organisms from exposure to extreme temperature and UV (Kemppinen *et al*., 2024).

Through spring and summer, when both temperature and UV radiation tend to increase, community visible reflectance decreased, indicating that butterfly communities exhibit darker colouration later in the season. Correspondingly, community visible reflectance also decreased as temperature increased. We expect that this darker colouration is predominantly driven by elevated UV rather than higher temperatures *per se*. Indeed, several studies have found that communities can become darker in hotter environments when UV increases (Bishop *et al*., 2016; Law *et al*., 2020). Although we are unaware of studies that address how colouration varies across communities through a year, this result contrasts with seasonal changes in *intra*specific reflectance patterns, as multivoltine species often have lighter-coloured generations in warmer parts of the year (Hoffmann, 1973). Our community level results also contrast with previous work showing that darker butterflies were associated with lower temperatures over microclimatic (Xing *et al*., 2016) and geographic scales (Munro *et al*., 2019; Kang *et al*., 2021). While darker colouration is expected from the UV-protection perspective (Bastide *et al*., 2014; Reguera *et al*., 2014; Delhey, 2017; Delhey, 2018; Delhey, 2019), we hypothesised that NIR reflectance would increase through the year to protect from the negative effects of high temperature in the summer. While we did not find that community NIR reflectance increased, we did find that it remained stable through the year, despite the strong correlation between NIR and visible reflectance across species, suggesting that different selection pressures could be acting on each reflectance type. The difference in magnitude between visible and NIR responses to temperature resembles a pattern observed in ant communities, which were darker in environments with higher temperatures and UV radiation, likely due to increased pigmentation for UV protection (Law *et al*., 2020), consistent with a trade-off between thermoregulation and UV protection. Given that visible reflectance is influenced by pigments required for protection against UV radiation (Alla *et al*., 2009), butterflies in warmer and high UV radiation environments would typically have lower visible reflectance. Conversely, to regulate their body temperatures to avoid overheating, butterflies would benefit from higher reflectance. Since NIR reflectance is likely due to structural colour rather than pigments (Tada *et al*., 1999; Kinoshita *et al*., 2002; Stavenga, 2021), it could respond in the opposing direction to visible reflectance to aid in thermoregulation. In our study, this trade-off might explain the difference in magnitudes of the relationships between visible and NIR reflectance as temperatures increase. Differences in visible and NIR reflectance patterns were also observed in Australian butterflies, which found that the relationship between climate and reflectance was stronger for NIR than visible wavelengths (Munro *et al*., 2019). Additionally, this has been observed in Australian birds, where birds living in hotter conditions have higher NIR reflectance, independent of their visible reflectance (Medina *et al*., 2018).

Our species-level analysis showed the opposite trend to community level trends, where butterfly species were instead lighter through spring and summer and at higher temperatures. While this may seem counterintuitive, we believe this to be an example of Simpson’s paradox (Wagner, 1982), where within-group trends are reversed when data are pooled, as the community-level models do not account for phylogeny and may therefore conflate within-group and between-group variation. In this case, species within specific butterfly clades get lighter throughout the year, but the lighter clades tend to come out earlier. Indeed, both visible and NIR reflectance exhibited strong phylogenetic signal, indicating that closely related species share similar colouration phenotypes, which may not be surprising given that butterfly families are often associated with particular colours (e.g. Pieridae contain the “Whites” and Lycaenidae contain the “Blues”). At the community level, earlier in the year, communities are dominated by lighter-coloured species from the Pieridae family, whereas darker-coloured species from the Nymphalidae family are more frequent later in the year. Once phylogeny is accounted for, however, the within-family trends reveal that darker individuals of each family are more abundant earlier in the year, while the lighter individuals of the same family are more abundant later in the year. These opposing trends could be driven by differences in thermoregulatory strategies of particular butterfly clades (Alamo *et al*., 2024). For example, butterflies in the Pieridae family employ a distinct behavioural mechanism for heat gain, which allows them to raise their body temperatures more than 11 - 12°C above ambient by reflecting sunlight down off their light wings onto their dark thoraxes (Kingsolver, 1985b). This thermoregulation strategy enables them to be active at lower ambient temperatures, allowing butterflies in the Pieridae family to emerge earlier in the season, despite being light coloured butterflies. The relatively stable community NIR reflectance through the year, in contrast to the decreasing community visible reflectance, shows that NIR reflectance does not passively mirror visible reflectance patterns despite visible and NIR reflectance being strongly coupled. Indeed, we found that species with higher NIR to visible reflectance ratios were more abundant later in the year (Figure S1). Together, these results suggest a trade-off between thermoregulation and protection against UV radiation. Visible reflectance depends on the amount of light absorbed by pigments such as melanin, which provides UV protection (Alla *et al*., 2009; Kang *et al*., 2021) while largely reflecting NIR wavelengths (Vogel *et al*., 1991; Kollias, 1995; Wistrand *et al*., 1997). On the other hand, structural colour, which is frequently found in butterflies due to the structure of the scales on their wings, can generate a broad range of spectral patterns with multiple peaks in various regions of the spectrum (Tada *et al*., 1999; Kinoshita *et al*., 2002; Stavenga, 2021). Such variation allows some decoupling of visible and NIR reflectances, but constraints do still cause strong correlations between the two (Munro *et al*., 2019; Kang *et al*., 2021).

There was no consistent trend in visible or NIR reflectance between our two time periods (2004/5 to 2017) at the community- or species-level, with only moderate support for a slight decrease in visible reflectance. Overall, the absence of a strong trend suggests that either the pace of climate change in the Guadarrama mountain range has not been sufficient to drive detectable shifts in colouration of butterfly communities over such a short time period, or that opposing selective pressures maybe be acting to constrain changes in reflectance. This contrasts broader-scale studies showing that climate change has driven changes in colouration, where darker coloured insect forms were less frequent when temperatures increased as a result of climate change (Zeuss *et al*., 2014; Clusella-Trullas and Nielsen, 2020).

We observed a non-linear relationship between both visible and NIR reflectance with UV radiation across communities. Visible reflectance of butterfly communities decreased as UV radiation increased from low to intermediate levels but increased at higher UV levels. These changes were much less pronounced for NIR reflectance, which showed a more linear change as UV increased. This difference in NIR reflectance compared to visible reflectance is consistent with the reflectance patterns through the year and with temperature, again suggesting that visible reflectance is more sensitive to UV radiation due to the role of melanin in protecting against UV radiation. At the species-level, UV radiation had no significant effect on either visible or NIR reflectance. We expect that these complex patterns of reflectance mirror the complexity of UV radiation varying across multiple spatiotemporal gradients, sometimes non-linearly (e.g. through the year).

Future work could build upon our analyses by accounting for within-species and body-region specific variation in reflectance. In this study, we used mean reflectance values averaged across six body regions per species. However, incorporating the relative surface area and function of each body region would enable more precise estimates of whole-body reflectance. For example, wing bases, thoraces, and distal wing regions differ in their roles in thermoregulation and display (Munro *et al*., 2019). Similarly, previous research found that basal wing regions showed a stronger relationship between reflectance and temperature than thoracic regions, suggesting region-specific thermal adaptation (Kang *et al*., 2021). To better understand how vegetation cover shapes these patterns, we propose using microclimate models that can incorporate vegetation (Maclean *et al*., 2019; Kearney and Porter, 2020) to determine whether the observed reflectance patterns across altitudes were related to differences in vegetation cover. Further integration of NIR and visible reflectance into biophysical models could improve mechanistic predictions of species’ thermal balance under climate change (Maclean *et al*., 2019; Kearney and Porter, 2020; Briscoe *et al*., 2023). Incorporating reflectance into a butterfly-specific model of operative body temperature would allow us to quantify how differences in pigmentation and structural colouration influence thermal exposure across thermal gradients in space and time. Additionally, linking reflectance data to species traits such as habitat use, body size and activity time could clarify how colour-mediated thermal strategies vary across ecological contexts.

## 5 Conclusions

By combining data from an extensive butterfly monitoring program and publicly available colouration data with climate data modelled at fine spatial scales, we show strong patterns in the visible and near-infrared (NIR) reflectance of butterfly species and communities in both space and time. Importantly, our results show that NIR reflectance does not simply mirror visible reflectance, despite strong correlation between the two. This partial decoupling highlights that NIR reflectance is an underexplored functional trait that can respond independently from visible reflectance over spatiotemporal and climatic gradients. Understanding how NIR reflectance varies across taxa is therefore critical for predicting how biodiversity will respond to future climate change. Furthermore, we highlight the vital importance for understanding evolutionary context when studying the ecology of extant organisms. Without accounting for phylogeny, we would incorrectly assume that daker butterflies dominated the summer months, when in fact, the opposite pattern is true. It is therefore imperative for ecologists to use phylogenetic data when understanding the complex relationships between organisms, their traits and the environment.

## References

1. Alamo M., García-Barros E. & Romo H. (2024). Adult thermoregulatory behaviour does not provide, by itself, an adaptive explanation for the reflectance–climate relationship (Bogert’s pattern) in Iberian butterflies. Ecological Entomology, 49(1), 77–90.

2. Alla S. K., Clark J. F. & Beyette F. R. Signal processing system to extract serum bilirubin concentration from diffuse reflectance spectrum of human skin. Proceedings of 2009 Annual International Conference of the IEEE Engineering in Medicine and Biology Society, 2009. pp. 1290–1293. (IEEE) 10.1109/IEMBS.2009.5333236

3. Álvarez H. A., Walker E., Mingarro M., Ursul G., Cancela J. P., Bassett L. & Wilson R. J. (2024). Heterogeneity in habitat and microclimate delay butterfly community tracking of climate change over an elevation gradient. Biological Conservation, 289, 110389.

4. Angilletta Jr M. J. (2009). Thermal adaptation: a theoretical and empirical synthesis.

5. Bastide H., Yassin A., Johanning E. J. & Pool J. E. (2014). Pigmentation in Drosophila melanogaster reaches its maximum in Ethiopia and correlates most strongly with ultra-violet radiation in sub-Saharan Africa. BMC evolutionary biology, 14(1), 179.

6. Bates D., Mächler M., Bolker B. & Walker S. (2014). Fitting linear mixed-effects models using lme4. arXiv preprint. 10.48550/arXiv.1406.5823

7. Beldade P. & Brakefield P. M. (2002). The genetics and evo–devo of butterfly wing patterns. Nature Reviews Genetics, 3(6), 442–452. 10.1038/nrg818

8. Bellard C., Bertelsmeier C., Leadley P., Thuiller W. & Courchamp F. (2012). Impacts of climate change on the future of biodiversity. Ecology letters, 15(4), 365–377. 10.1111/j.1461-0248.2011.01736.x

9. Bishop T. R., Robertson M. P., Gibb H., Van Rensburg B. J., Braschler B., Chown S. L., Foord S. H., Munyai T. C., Okey I. & Tshivhandekano P. G. (2016). Ant assemblages have darker and larger members in cold environments. Global Ecology and Biogeography, 25(12), 1489–1499. 10.1111/geb.12516

10. Bogert C. M. (1949). Thermoregulation in reptiles, a factor in evolution. Evolution, 3(3), 195–211. 10.2307/2405558

11. Bridle J., Balmford A., Durant S. M., Gregory R. D., Pearson R. & Purvis A. (2025). How should we bend the curve of biodiversity loss to build a just and sustainable future? Page 20230205. The Royal Society.

12. Brindle M., Ferguson-Gow H., Williamson J., Thomsen R. & Sommer V. (2023). The evolution of masturbation is associated with postcopulatory selection and pathogen avoidance in primates. Proceedings of the Royal Society B, 290(2000), 20230061.

13. Briscoe N. J., Morris S. D., Mathewson P. D., Buckley L. B., Jusup M., Levy O., Maclean I. M., Pincebourde S., Riddell E. A. & Roberts J. A. (2023). Mechanistic forecasts of species responses to climate change: the promise of biophysical ecology. Global Change Biology, 29(6), 1451–1470. 10.1111/gcb.16557

14. Bürkner P.-C. (2021). Bayesian Item Response Modeling in R with brms and Stan. Journal of Statistical Software, 100(5), 1–54. 10.18637/jss.v100.i05

15. Calosi P., Bilton D. T. & Spicer J. I. (2008). Thermal tolerance, acclimatory capacity and vulnerability to global climate change. Biology letters, 4(1), 99–102. 10.1098/rsbl.2007.0408

16. Chevin L.-M. & Bridle J. (2025). Impacts of limits to adaptation on population and community persistence in a changing environment. Philosophical Transactions B, 380(1917), 20230322.

17. Clusella-Trullas S. & Nielsen M. (2020). The evolution of insect body coloration under changing climates. Current Opinion in Insect Science, 41, 25–32. 10.1016/j.cois.2020.05.007

18. Cuthill I. C., Allen W. L., Arbuckle K., Caspers B., Chaplin G., Hauber M. E., Hill G. E., Jablonski N. G., Jiggins C. D. & Kelber A. (2017). The biology of color. Science, 357(6350), eaan0221. 10.1126/science.aan0221

19. de Souza A. R., Mayorquin A. Z. & Sarmiento C. E. (2020). Paper wasps are darker at high elevation. Journal of Thermal Biology, 89, 102535. 10.1016/j.jtherbio.2020.102535

20. Delhey K. (2017). Gloger’s rule. Current Biology, 27(14), R689–R691.

21. Delhey K. (2018). Darker where cold and wet: Australian birds follow their own version of Gloger’s rule. Ecography, 41(4), 673–683. 10.1111/ecog.03040

22. Delhey K. (2019). A review of Gloger’s rule, an ecogeographical rule of colour: Definitions, interpretations and evidence. Biological Reviews, 94(4), 1294–1316. 10.1111/brv.12503

23. Deutsch C. A., Tewksbury J. J., Huey R. B., Sheldon K. S., Ghalambor C. K., Haak D. C. & Martin P. R. (2008). Impacts of climate warming on terrestrial ectotherms across latitude. Proceedings of the National Academy of Sciences, 105(18), 6668–6672.

24. Gautam S. & Kunte K. (2020). Adaptive plasticity in wing melanisation of a montane butterfly across a Himalayan elevational gradient. Ecological Entomology, 45(6), 1272–1283. 10.1111/een.12911

25. Goded M., Ursul G., Baz A. & Wilson R. J. (2024). Changes to butterfly phenology versus elevation range after four decades of warming in the mountains of central Spain. Journal of Insect Conservation, 1–15. 10.1007/s10841-024-00561-8

26. Hersbach H., Bell B., Berrisford P., Hirahara S., Horányi A., Muñoz-Sabater J., Nicolas J., Peubey C., Radu R. & Schepers D. (2020). The ERA5 global reanalysis. Quarterly journal of the royal meteorological society, 146(730), 1999–2049.

27. Hoffmann R. J. (1973). Environmental control of seasonal variation in the butterfly Colias eurytheme. I. Adaptive aspects of a photoperiodic response. Evolution, 387–397. 10.2307/2407302

28. Hollister J. W., Shah T., Nowosad J., Robitaille A. L., Beck M. W. & Johnson M. (2023). elevatr: Access Elevation Data from Various APIs, R package version 0.99.1. GitHub: Available at https://github.com/usepa/elevatr/

29. Illán J. G., Gutierrez D., Diez S. B. & Wilson R. J. (2012). Elevational trends in butterfly phenology: implications for species responses to climate change. Ecological Entomology, 37(2), 134–144. 10.1111/j.1365-2311.2012.01345.x

30. Illán J. G., Gutiérrez D. & Wilson R. J. (2010). The contributions of topoclimate and land cover to species distributions and abundance: fine-resolution tests for a mountain butterfly fauna. Global Ecology and Biogeography, 19(2), 159–173. 10.1111/j.1466-8238.2009.00507.x

31. Kalmus H. (1941). The resistance to desiccation of Drosophila mutants affecting body colour. Proceedings of the Royal Society of London. Series B-Biological Sciences, 130(859), 185–201.

32. Kang C., Im S., Lee W. Y., Choi Y., Stuart-Fox D. & Huertas B. (2021). Climate predicts both visible and near-infrared reflectance in butterflies. Ecology letters, 24(9), 1869–1879. 10.1111/ele.13821

33. Kearney M. R. & Porter W. P. (2020). NicheMapR–an R package for biophysical modelling: the ectotherm and dynamic energy budget models. Ecography, 43(1), 85–96. 10.1111/ecog.04680

34. Kemppinen J., Lembrechts J. J., Van Meerbeek K., Carnicer J., Chardon N. I., Kardol P., Lenoir J., Liu D., Maclean I. & Pergl J. (2024). Microclimate, an important part of ecology and biogeography. Global Ecology and Biogeography, 33(6), e13834.

35. Kettlewell H. B. D. (1955). Selection experiments on industrial melanism in the Lepidoptera. Heredity, 9(3), 323–342.

36. King K. J. & Sinclair B. J. (2015). Water loss in tree weta (Hemideina): adaptation to the montane environment and a test of the melanisation–desiccation resistance hypothesis. The Journal of Experimental Biology, 218(13), 1995–2004.

37. Kingsolver J. G. (1985a). Butterfly thermoregulation: organismic mechanisms and population consequences. Journal of Research on the Lepidoptera, 24(1), 1–20.

38. Kingsolver J. G. (1985b). Thermal ecology of Pieris butterflies (Lepidoptera: Pieridae): a new mechanism of behavioral thermoregulation. Oecologia, 66, 540–545. 10.1007/BF00379347

39. Kingsolver J. G. (1988). Thermoregulation, flight, and the evolution of wing pattern in pierid butterflies: the topography of adaptive landscapes. American Zoologist, 28(3), 899–912. 10.1093/icb/28.3.899

40. Kinoshita S., Yoshioka S. & Kawagoe K. (2002). Mechanisms of structural colour in the Morpho butterfly: cooperation of regularity and irregularity in an iridescent scale. Proceedings of the Royal Society of London. Series B: Biological Sciences, 269(1499), 1417–1421. 10.1098/rspb.2002.2019

41. Klinges D. H., Duffy J. P., Kearney M. R. & Maclean I. M. (2022). mcera5: Driving microclimate models with ERA5 global gridded climate data. Methods in Ecology and Evolution, 13(7), 1402–1411. 10.1111/2041-210X.13877

42. Kollias N. (1995). The Spectroscopy of Human Melanin Pigmentation.

43. Law S. J., Bishop T. R., Eggleton P., Griffiths H., Ashton L. & Parr C. (2020). Darker ants dominate the canopy: testing macroecological hypotheses for patterns in colour along a microclimatic gradient. Journal of Animal Ecology, 89(2), 347–359. 10.1111/1365-2656.13110

44. Luo D.-G., Yue W. W., Ala-Laurila P. & Yau K.-W. (2011). Activation of visual pigments by light and heat. Science, 332(6035), 1307–1312. 10.1126/science.1200172

45. Maclean I. M., Mosedale J. R. & Bennie J. J. (2019). Microclima: An r package for modelling meso-and microclimate. Methods in Ecology and Evolution, 10(2), 280–290. 10.1111/2041-210X.13093

46. Maclean I. M. D. & Mosedale J. R. (2025). mesoclim. https://github.com/ilyamaclean/mesoclim.

47. Majerus M. E. (1998) ’Melanism: evolution in action.’ (Oxford University Press)

48. Medina I., Newton E., Kearney M. R., Mulder R. A., Porter W. P. & Stuart-Fox D. (2018). Reflection of near-infrared light confers thermal protection in birds. Nature Communications, 9(1), 3610. 10.1038/s41467-018-05898-8

49. Mingarro M., Cancela J. P., BurÓn-Ugarte A., García-Barros E., Munguira M. L., Romo H. & Wilson R. J. (2021). Butterfly communities track climatic variation over space but not time in the Iberian Peninsula. Insect Conservation and Diversity, 14(5), 647–660. 10.1111/icad.12498

50. Munro J. T., Medina I., Walker K., Moussalli A., Kearney M. R., Dyer A. G., Garcia J., Rankin K. J. & Stuart-Fox D. (2019). Climate is a strong predictor of near-infrared reflectance but a poor predictor of colour in butterflies. Proceedings of the Royal Society B, 286(1898), 20190234. 10.1098/rspb.2019.0234

51. New M., Lister D., Hulme M. & Makin I. (2002). A high-resolution data set of surface climate over global land areas. Climate research, 21(1), 1–25.

52. Parkash R., Ramniwas S., Rajpurohit S. & Sharma V. (2008). Variations in body melanization impact desiccation resistance in Drosophila immigrans from Western Himalayas. Journal of Zoology, 276(2), 219–227.

53. Parkash R., Singh S. & Ramniwas S. (2009). Seasonal changes in humidity level in the tropics impact body color polymorphism and desiccation resistance in Drosophila jambulina—Evidence for melanism-desiccation hypothesis. Journal of insect physiology, 55(4), 358–368.

54. Parmesan C., Morecroft M. D., Trisurat Y., Adrian R., Anshari G. Z., Arneth A., Gao Q., Gonzalez P., Harris R., Price J., Stevens N. & Talukdarr G. H. (2022) Chapter 2: Terrestrial and Freshwater Ecosystems and Their Services. IPCC, https://www.ipcc.ch/report/ar6/wg2/chapter/chapter-2/. 10.1017/9781009325844.004

55. Pinsky M. L., Eikeset A. M., McCauley D. J., Payne J. L. & Sunday J. M. (2019). Greater vulnerability to warming of marine versus terrestrial ectotherms. Nature, 569(7754), 108–111.

56. Pollard E. & Yates T. J. (1993) ’Monitoring butterflies for ecology and conservation: the British butterfly monitoring scheme.’ (Springer Science & Business Media)

57. Pottier P., Kearney M. R., Wu N. C., Gunderson A. R., Rej J. E., Rivera-Villanueva A. N., Pollo P., Burke S., Drobniak S. M. & Nakagawa S. (2025). Vulnerability of amphibians to global warming. Nature, 1–8.

58. R Core Team (2024). *R: A Language and Environment for Statistical Computing*, R version 4.3.3 (2024-02-29 ucrt). R Foundation for Statistical Computing, Vienna, Austria. Available at https://www.R-project.org/

59. Reguera S., Zamora-Camacho F. J. & Moreno-Rueda G. (2014). The lizard Psammodromus algirus (Squamata: Lacertidae) is darker at high altitudes. Biological Journal of the Linnean Society, 112(1), 132–141. 10.1111/bij.12250

60. Rockstrom J., Steffen W. & Noone K. (2009). A safe operating space for humanity. Nature, 461(7263), 472–475.

61. Román-Palacios C. & Wiens J. J. (2020). Recent responses to climate change reveal the drivers of species extinction and survival. Proceedings of the National Academy of Sciences, 117(8), 4211–4217. 10.1073/pnas.1913007117

62. Schirmer S. C., Gawryszewski F. M., Cardoso M. Z. & Pessoa D. M. A. (2023). Melanism and color saturation of butterfly assemblages: A comparison between a tropical rainforest and a xeric white forest. Frontiers in Ecology and Evolution, 11, 932755.

63. Stavenga D. G. (2021). The wing scales of the mother-of-pearl butterfly, Protogoniomorpha parhassus, are thin film reflectors causing strong iridescence and polarization. Journal of Experimental Biology, 224(15), jeb242983. 10.1242/jeb.242983

64. Stuart-Fox D., Newton E. & Clusella-Trullas S. (2017). Thermal consequences of colour and near-infrared reflectance. Philosophical Transactions of the Royal Society B: Biological Sciences, 372(1724), 20160345. 10.1098/rstb.2016.0345

65. Tada H., Mann S. E., Miaoulis I. N. & Wong P. Y. (1999). Effects of a butterfly scale microstructure on the iridescent color observed at different angles. Optics Express, 5(4), 87–92.

66. True J. R. (2003). Insect melanism: the molecules matter. Trends in ecology & evolution, 18(12), 640–647.

67. Tsai C.-C., Childers R. A., Nan Shi N., Ren C., Pelaez J. N., Bernard G. D., Pierce N. E. & Yu N. (2020). Physical and behavioral adaptations to prevent overheating of the living wings of butterflies. Nature communications, 11(1), 551. 10.1038/s41467-020-14408-8

68. Urban M. C. (2015). Accelerating extinction risk from climate change. Science, 348(6234), 571–573.

69. Vogel A., Dlugos C., Nuffer R. & Birngruber R. (1991). Optical properties of human sclera, and their consequences for transscleral laser applications. Lasers in surgery and medicine, 11(4), 331–340. 10.1002/lsm.1900110404

70. Wagner C. H. (1982). Simpson’s paradox in real life. The American Statistician, 36(1), 46–48.

71. Wiemers M. (2024). Email to Martin Wiemers.

72. Wiemers M., Chazot N., Wheat C. W., Schweiger O. & Wahlberg N. (2020). A complete time-calibrated multi-gene phylogeny of the European butterflies. ZooKeys, 938, 97. 10.3897/zookeys.938.50878

73. Wiens J. J. (2016). Climate-related local extinctions are already widespread among plant and animal species. PLoS biology, 14(12), e2001104. 10.1371/journal.pbio.2001104

74. Wiernasz D. C. (1995). Male choice on the basis of female melanin pattern in Pieris butterflies. Animal Behaviour, 49(1), 45–51.

75. Williamson J., Lu M., Camus M. F., Gregory R. D., Maclean I., Rocha J. C., Saastamoinen M., Wilson R. J., Bridle J. & Pigot A. L. (2025). Clustered warming tolerances and the nonlinear risks of biodiversity loss on a warming planet. Philosophical Transactions of the Royal Society B: Biological Sciences, 380(1917).

76. Williamson J., Teh E., Jucker T., Brindle M., Bush E., Chung A. Y., Parrett J., Lewis O. T., Rossiter S. J. & Slade E. M. (2022). Local-scale temperature gradients driven by human disturbance shape the physiological and morphological traits of dung beetle communities in a Bornean oil palm–forest mosaic. Functional Ecology, 36(7), 1655–1667. 10.1111/1365-2435.14062

77. Wilson R. J., Gutiérrez D., Gutiérrez J., Martínez D., Agudo R. & Monserrat V. J. (2005). Changes to the elevational limits and extent of species ranges associated with climate change. Ecology letters, 8(11), 1138–1146. 10.1111/j.1461-0248.2005.00824.x

78. Wilson R. J., Gutierrez D., Gutierrez J. & Monserrat V. J. (2007). An elevational shift in butterfly species richness and composition accompanying recent climate change. Global Change Biology, 13(9), 1873–1887. 10.1111/j.1365-2486.2007.01418.x

79. Wistrand P. J., Stjernschantz J. & Olsson K. (1997). The incidence and time-course of latanoprost-induced iridial pigmentation as a function of eye color. Survey of ophthalmology, 41, S129–S138. 10.1016/S0039-6257(97)80020-3

80. Xing S., Bonebrake T. C., Tang C. C., Pickett E. J., Cheng W., Greenspan S. E., Williams S. E. & Scheffers B. R. (2016). Cool habitats support darker and bigger butterflies in Australian tropical forests. Ecology and Evolution, 6(22), 8062–8074. 10.1002/ece3.2464

81. Zeuss D., Brandl R., Brändle M., Rahbek C. & Brunzel S. (2014). Global warming favours light-coloured insects in Europe. Nature Communications, 5(1), 3874. 10.1038/ncomms4874

