## Supplementary material for "Temperature and Ultraviolet Radiation Drive Divergent Visible and Near-Infrared Reflectance Patterns in Butterflies"

**Supporting Information** for paper "Temperature and Ultraviolet Radiation Drive Divergent Visible and Near-Infrared Reflectance Patterns in Butterflies"

### Supplementary methods

*Residuals analysis*

To explore the relationship between visible and near-infrared (NIR) reflectance across butterfly species in our study system, we fit a linear model using NIR reflectance as the response variable and visible reflectance as the predictor variable. This model provided an estimate of the overall relationship between visible and NIR reflectance for butterfly species in our study system. The residuals from this model represent the difference between the observed NIR reflectance and the expected NIR reflectance predicted by the model based on visible reflectance. Positive residuals indicate higher NIR reflectance than expected, while negative residuals indicate lower NIR reflectance than expected. Following this, we calculated the community-weighted mean of the residuals of the previous model for each transect. To understand whether differences in the patterns of each reflectance type are due to the NIR to visible reflectance ratios of species, we fit a linear mixed-effects model with community-weighted residuals as the response variable, altitude and day of the year as continuous fixed effects, year as a categorical fixed effect and transect location as a random effect.

### Supplementary results

*Species-level analysis*

**Table S1.** Results of species-level analysis.

| Predictor | Reflectance | Mean | Lower 95 % BCI | Lower 80 % BCI | Upper 95 % BCI | Upper 80 % BCI | Support |
|---|---|---|---|---|---|---|---|
| Altitude (m) | Visible | -1.34 | -2.89 | -2.34 | 0.22 | -0.33 | Moderate |
| Altitude (m) | NIR | -2.56 | -4.79 | -4.01 | -0.33 | -1.11 | Strong |
| Day of the year | Visible | 1.38 | -0.27 | 0.31 | 3.03 | 2.44 | Moderate |
| Day of the year | NIR | 2.66 | 0.33 | 1.14 | 4.98 | 4.16 | Strong |
| Time period | Visible | -0.87 | -2.10 | -1.66 | 0.37 | -0.07 | Moderate |
| Time period | NIR | -0.93 | -2.87 | -2.17 | 0.99 | 0.32 | None |
| Temperature (°C) | Visible | 1.27 | -0.28 | 0.26 | 2.81 | 2.27 | Moderate |
| Temperature (°C) | NIR | 2.52 | 0.33 | 1.10 | 4.71 | 3.94 | Strong |
| UV Radiation ($W/m^2$) | Visible | -0.39 | -1.98 | -1.42 | 1.19 | 0.64 | None |
| UV Radiation ($W/m^2$) | NIR | -0.70 | -2.98 | -2.17 | 1.56 | 0.78 | None |

*Residuals analysis*

The best fitting model for our residuals analysis included day of the year, which significantly improved model fit ($x_1^2$ = 318.94, p < 0.001, LRT compared to model without day of the year; Figure S1). Through the year, butterfly species had higher near-infrared to visible reflectance (Figure S1).

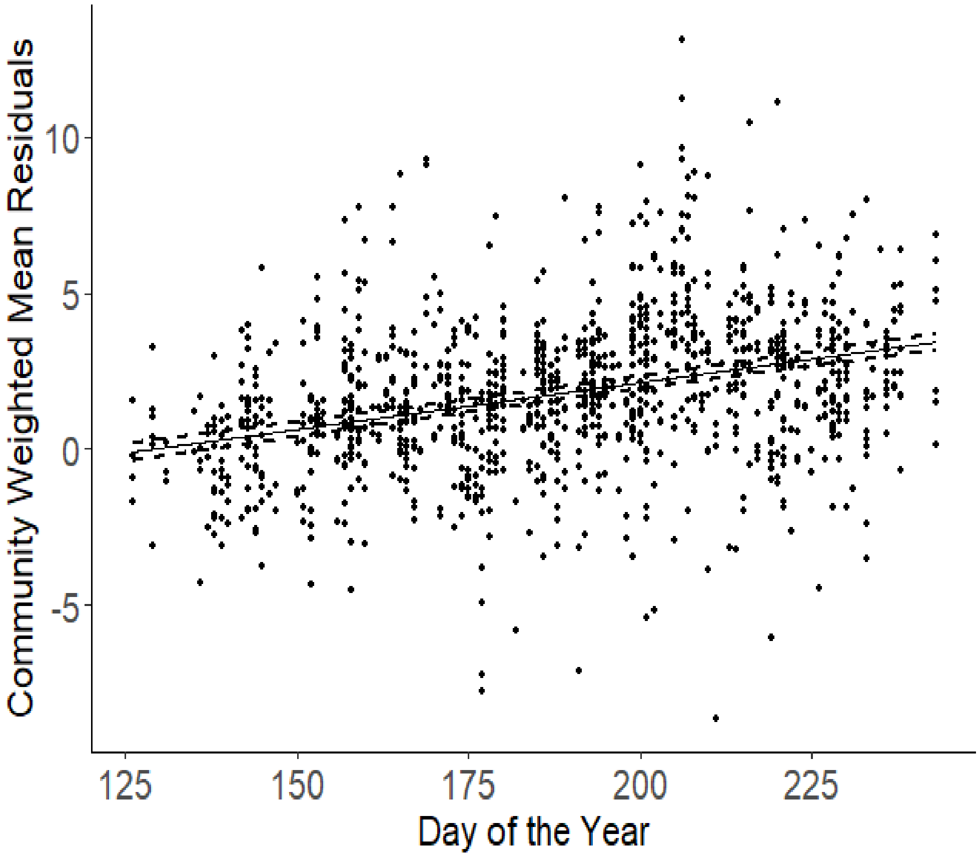


**Figure S1. Community-weighted mean residuals through the year.** Dashed lines denote 95 % confidence intervals.
